# The conserved N-terminal SANT1-binding domain (SBD) of EZH2 Regulates PRC2 Activity

**DOI:** 10.1101/2025.02.04.636462

**Authors:** Agata L. Patriotis, Yadira Soto-Feliciano, Douglas W. Barrows, Laiba Khan, Marylene Leboeuf, Peder J. Lund, Matthew R Marunde, Annaelle Djomo, Michael-Christopher Keogh, Thomas S. Carroll, Benjamin A. Garcia, Alexey A. Soshnev, C. David Allis

## Abstract

Polycomb group proteins maintain gene expression patterns established during early development, with Polycomb Repressive Complex 2 (PRC2) methyltransferase a key regulator of cell differentiation, identity and plasticity. Consequently, extensive somatic mutations in PRC2, including gain- or loss- of function (GOF or LOF), are observed in human cancers. The regulation of chromatin structure by PRC2 is critically dependent on its EZH2 (Enhancer of Zeste Homolog 2) subunit, which catalyzes the methylation of histone H3 lysine 27 (H3K27). Recent structural studies of PRC2 revealed extensive conformational changes in the non-catalytic EZH2 N-terminal SANT-Binding Domain (SBD) during PRC2 activation, though the functional significance remains unclear. Here, we investigate how the SBD regulates PRC2 function. The domain is highly conserved in metazoans, dispensable for PRC2 assembly and chromatin localization, yet required for genome-wide histone H3K27 methylation. Further, we show that an intact SBD is necessary for the proliferation of EZH2- addicted lymphomas, and its deletion in the presence of *EZH2* GOF mutations inhibits cancer cell growth. These observations provide new insights to the regulation of PRC2 activity in normal development and malignancy.

## INTRODUCTION

The Polycomb Group proteins modify histones in chromatin, allowing for temporal and cell type-specific control of transcriptional programs during development. Dysregulation of these silencing mechanisms alters gene expression programs, leading to cancer and developmental disorders ^1–6^.

Polycomb Repressive Complex 2 (PRC2) methylates histone H3 lysine 27 (H3K27), with di- and tri-methylation (H3K27me2 and H3K27me3) localized to facultative heterochromatin: nuclear regions often encompassing developmentally repressed genes ^7–10^. The functional core of PRC2 is comprised of four proteins: three invariant non-enzymatic subunits (Embryonic Ectoderm Development (EED), Suppressor of Zeste 12 (SUZ12), and histone binding protein RBBP4/7 (also known as RbAp46/48)), and one of the two SET domain-containing methyltransferases Enhancer of Zeste Homolog 1 or 2 (EZH1 or 2) ^10^.

Multiple molecular mechanisms regulate PRC2 activity. In vertebrates, broad H3K27me3 domains are initiated when PRC2 is first recruited to strong nucleation sites enriched in CpG islands ^11^. This is followed by allosteric enzyme activation, successively leading to proximal spreading, distal nucleation, and further spreading by long-range chromatin interactions. PRC2 enzyme activity is further amplified by positive feedback from its terminal enzymatic product, H3K27me3 ^12,13^. Here the aromatic cage of the EED subunit specifically binds this post-translational modification (PTM), leading to allosteric stimulation of EZH2 methyltransferase activity, promoting the establishment of H3K27me3 across broad genomic regions ^13^. Of note, incorporation of EZH2 into PRC2 confers higher methyltransferase activity compared with EZH1 ^14^. Recently, a series of elegant biochemical and genetic studies have identified PRC2.1 and PRC2.2 complexes based on their distinct accessory proteins and associated regulatory activity. PRC2.1 is associated with Polycomb-like (PCL) proteins, including MTF2, and additional co-factors PALI1 or EPOP. Together these direct the complex to unmethylated CpG islands and bivalent promoters ^15^. Methylation of PALI1 then allosterically activates PRC2 for processive catalysis ^16^. PRC2.2, in contrast, is associated with JARID2 and AEBP2, with the former providing both regulatory inputs via allosteric activation and genome targeting to mono-ubiquitylated H2A lysine 119 (H2AK119Ub). AEBP2 is involved in PRC2 targeting via its DNA-binding zinc finger ^17^. Many PRC2 subunits, including specific accessory proteins and EZH1/2 subunits, show developmentally restricted expression profiles ^18^, suggesting that the establishment and maintenance of H3K27 methylation relies on distinct and diverse biochemical mechanisms (Sanulli et al. 2015; Boyer et al. 2006; Oktaba et al. 2008).

The function of the PRC2 complex is often dysregulated in cancer. Loss-of-function (LOF) mutations and deletions of *EZH2, EED*, and *SUZ12* genes are found in T-cell acute lymphoblastic leukemias (T-ALL) and malignant peripheral nerve sheath tumors (MPNSTs) ^19,20^. MPNSTs lacking EED or SUZ12 derepress known Polycomb-regulated target genes, including *Hox* genes, while re-introduction of the missing subunits restores repression and slows cellular proliferation ^19^. Overexpression of EZHIP, an inhibitory subunit of PRC2.2, and the resulting loss of H3K27me3, are frequent in pediatric ependymomas ^21^. Likewise, “oncohistone” H3 K27M which dominantly inhibits H3K27-specific methyltransferase activity, is a major driver in diffuse intrinsic pontine and midline gliomas ^22–24^. In contrast, EZH2 overexpression in cutaneous melanoma and cancers of the endometrium, prostate, and breast correlate with aggressiveness and poor prognosis ^25^. The “hotspot” mutations within the catalytic SET domain of EZH2 at tyrosine 641 (Y641) are frequent in lymphoma and melanoma ^4,26,27^. These gain-of-function (GOF) mutations increase the efficiency of H3K27me2ème3 conversion and simultaneously reduce the ability to deposit initial mono-methylation, and thus depend on a second, wild-type copy of EZH2 to catalyze the H3K27me0ème1 and H3K27me1ème2 reactions ^27^. Lymphomas carrying EZH2 GOF mutations often depend on PRC2/EZH2 for growth, and EZH2 inhibitor Tazemetostat is used in refractory follicular lymphoma with *EZH2* mutations ^28–30^.

The structure of EZH2 is complex, with several evolutionarily conserved domains (**Figure 1A**). This includes a C-terminal catalytic SET domain; a pre-SET cysteine-rich CXC domain necessary for efficient H3K27 tri-methylation and alternatively spliced in spermatocytes ^31^; two <u>S</u>wi3, <u>A</u>da2, <u>N</u>-CoR and <u>T</u>FIIIB (SANT) domains, of which SANT1 is implicated in recognition of an unmodified histone H4 tail ^32^; an autoregulatory <u>S</u>timulation-<u>R</u>esponsive <u>M</u>otif (SRM) involved in allosteric activation ^33,34^; and four shorter motifs within the N-terminal region, including SBD (SANT-Binding Domain), EBD (EED-Binding Domain), BAM (β-Addition Motif) and SAL (SET Activation Loop) ^33,35–38^. The N-terminal SBD was recently reported to undergo a conformational change upon allosteric activation of PRC2 ^39–42^, although its functional importance remains unclear.

**Figure 1.**
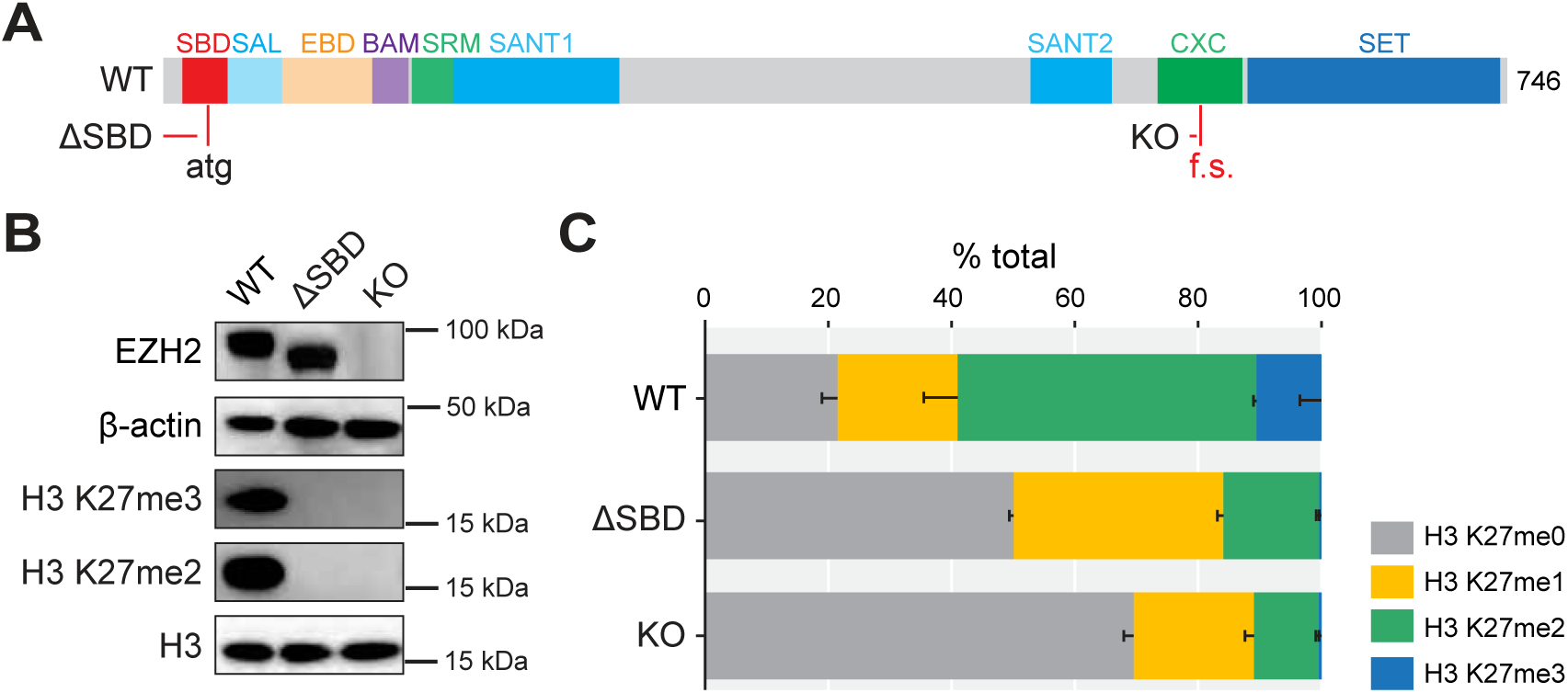
EZH2 SANT-binding domain (SBD) is required for PRC2-mediated H3K27 methylation. **A.** EZH2 contains several domains, including the poorly characterized N-terminal SBD (SANT1-binding domain). Location of mutations corresponding to ΔSBD and KO alleles is indicated: ΔSBD is an N-terminally truncated version of EZH2 lacking the first 22 amino acids; KO is a null mutation arising from a deletion leading to frameshift and premature stop codon in the CXC domain that does not accumulate EZH2 protein. **B.** Immunoblot of EZH2 and histone methylation in wild type mES cells (WT), isogenic homozygous ΔSBD EZH2 line (ΔSBD), and isogenic EZH2 knock-out line (KO). β-actin and histone H3 are loading controls. **C.** Tandem mass spectrometry (MS) analysis of H3K27 methylation states in acid-extracted histones from WT, ΔSBD and EZH2 KO mES cells. Error bars = SEM.

In this study, we interrogated the EZH2 SBD, hypothesizing that this highly conserved domain acts to regulate the H3K27 methylation activity of PRC2. To investigate SBD function, we disrupted the domain *in vitro* and *in vivo*, using both mouse embryonic stem cells (mESCs) and human lymphoma cell lines, followed by genome-wide analyses of H3K27 methylation, gene expression and cell proliferation. We show that SBD loss does not preclude PRC2 assembly yet dramatically reduces PRC2 methyltransferase activity. Further, an intact SBD is critical for the tumorigenic effect of the EZH2 Y641N GOF. Together, these results expand upon the complexity of PRC2 regulation *in vivo* and demonstrate that EZH2 catalytic activity is regulated by a distal, non-catalytic domain.

## RESULTS

### Conserved EZH2 SBD domain is necessary for H3K27 methylation by PRC2

Previous structural studies identified the EZH2 SBD as a bridge element between SANT1 and EED-binding domains ^40^. Spanning the first 32 amino acids of human EZH2, the region encompasses ten unstructured amino acids and the N-terminal portion of the first α-helix. To further investigate the significance of the EZH2 SBD, we first analyzed its sequence conservation between orthologs **(Supplemental Figure 1A)**. Several adjacent amino acids are conserved from *C. elegans* to human, suggesting potential functional importance ^43^. Conservation extends beyond the basic patches identified as likely DNA-binding motifs, suggesting additional functions ^40^. To study EZH2 SBD *in vivo*, we edited an mESC line to remove the first 22 amino acids of the endogenous protein (EZH2 ΔSBD, **Figure 1A**). To monitor resulting PRC2 enzymatic activity, we analyzed global histone H3K27 methylation levels by immunoblotting and bottom-up mass-spectrometry (MS). Removal of SBD leads to global depletion of H3K27me3 and H3K27me2, comparable to that observed on complete loss of EZH2 **(Figure 1B, C)**. These results demonstrate that the EZH2 SBD is indispensable for H3K27 methylation activity *in vivo*.

To further investigate the role of conserved sequences within the SBD, we replaced the endogenous domain from mouse *Ezh2* with orthologous regions from nematode (*C. elegans, Cel*), fruit fly (*D. melanogaster, Dme*) or turtle (*P. sinensis, Psi*). The resulting chimeric proteins were expressed in *Ezh2* KO mESCs to determine if they could restore methyltransferase activity and functionality. Here the SBD from *Dme* and *Psi*, but not that from *Cel*, restored H3K27me2 and H3K27me2 levels to EZH2-WT-like levels **(Supplemental Figure 1B)**. This was further corroborated by mESC lines expressing EZH2 chimeras in an assay for pluripotency maintenance **(Supplemental Figure 1C).** We induced differentiation by withdrawing Leukemia Inhibitory Factor (LIF) from the growth media of parental *Ezh2* KO mESCs and five transformed lines (EZH2 WT, EZH2 ΔSBD, EZH2 SBD-*Psi*, EZH2 SBD-*Dme*, and EZH2 SBD-*Cel*) followed by alkaline phosphatase (AP) activity staining ^44^. In this assay pluripotent ESCs are characterized by high AP activity ^45^. Loss of EZH2 results in a differentiation block, and maintenance of high AP activity, even upon LIF withdrawal ^31,46,47^ **(Supplemental Figure 1C).** Consistent with immunoblot analyses, only those EZH2 SBD constructs that restored H3K27 methylation (EZH2 WT, EZH2 SBD-*Psi*, and EZH2 SBD-*Dme*) restored mESC differentiation, demonstrating that the evolutionarily conserved SBD is necessary for appropriate PRC2 function.

### EZH2 SBD is essential for effective H3K27 methylation

Despite significant reduction, residual H3K27me2 and H3K27me3 were still detected by MS in EZH2 ΔSBD cells (**Figure 1C**). We thus performed reference-controlled Chromatin ImmunoPrecipitation followed by next-generation sequencing (ChIP-Rx) to quantitatively characterize the genomic distribution of these methyl states ^48^. While EZH2 ΔSBD line showed dramatically reduced H3K27 methylation across PRC2 targets, residual H3K27me2 signal accumulated at the EZH2 WT H3K27me3 peak regions **(Figure 2).** As the accumulation of H3K27me2 at sites where PRC2 complex recruitment is initiated precedes the formation of broad H3K27me3 domains ^11^, we hypothesized that SBD may contribute to EZH2 allosteric activation.

**Figure 2.**
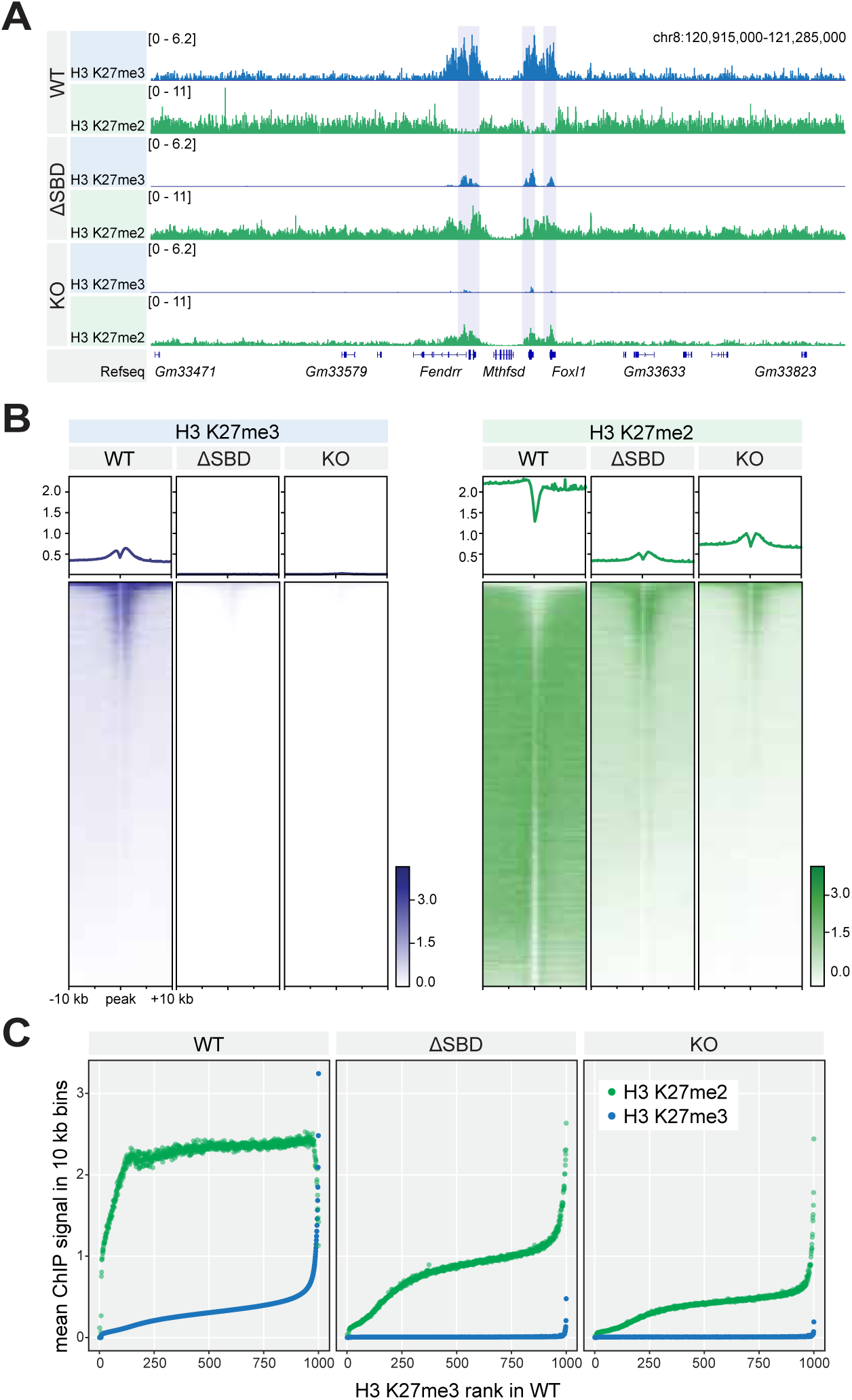
Partial deletion of EZH2 SBD domain leads to the global loss of H3K27me3 and redistribution of H3K27me2. **A.** Genome browser view of H3K27me3 (blue) and H3K27me2 (green) ChIP-Rx signal. Refseq annotations are shown below. Shaded boxes highlight retained H3K27me3 sites in EZH2-ΔSBD cells. **B.** Heatmaps of H3K27me3 (blue) and H3K27me2 (green) ChIP-Rx signal within a 20 kb window centered on transcription start sites (TSSs) in EZH2 WT mESCs. **C.** ChIP-Rx normalized reads per 10 kb bin for H3K27me3 (blue) and H3K27me2 (green) in parental (EZH2 WT), EZH2 SET KO and EZH2-ΔSBD mESCs show the replacement of H3K27me3 signal with H3K27me2 in both mutant backgrounds. 1 kb genomic tiles were ranked by H3K27me2 and H3K27me3 enrichment in EZH2 WT mESCs and grouped into 1,000 bins ordered by rank.

To investigate how SBD contributes to the formation of broad H3K27me3 domains, we compared our ChIP-Rx data to the dynamics of PRC2 complex recruitment to nucleation sites in the mESC model ^11^. PRC2 recruitment is followed by allosteric activation and proximal spreading, followed by distal nucleation and spreading via long-range chromatin interactions ^11^ The EED Y365A mutant limits allosteric activation of the PRC2 complex ^49^, and identifies sites of early PRC2 recruitment ^11^. Of note, H3K27me3 distribution in EZH2 ΔSBD cells bears remarkable similarity to PRC2 nucleation sites in EED Y365A cells ^9,11,50^ (**Supplemental Figure 2A**). In the EZH2 ΔSBD mESCs H3K27me3 persisted in distinct, narrow sites embedded within larger Polycomb domains, with EZH2 strictly localized to the same regions **(Supplemental Figure 2B,D)**. Further, as PRC2 nucleation sites are characterized by high density of CpG islands compared to broader H3K27me3 domains ^11,51^, we compared the H3K27me3 signal with CpG content in parental and EZH2 ΔSBD cell, and found overlapping enrichment of retained peaks with CpG islands in EZH2 ΔSBD cells **(Supplemental Figure 2C)**. Together, these findings suggest that EZH2 ΔSBD is effectively recruited to strong nucleation sites, but fails to attain the local methylation levels necessary to initiate the spreading of H3K27me3.

### EZH2 SBD is required for PRC2 enzymatic activity independent of complex composition stability or recruitment to chromatin

Structural studies of PRC2 complexed with a H3K27me3 nucleosome describe engagement of the EZH2 N-terminal region with the EZH2 SANT1 domain, EED and DNA ^40,52^. As such, we surmised that SBD loss may impair PRC2 activity by directly impacting complex composition, stability and/or recruitment to chromatin. To investigate this, we used co-immunoprecipitation (co-IP) via SUZ12 followed by MS in EZH2 WT, KO, and ΔSBD mESCs. Curiously, the results showed PRC2 assembly is independent of its methyltransferase subunit, and loss of SBD did not preclude EZH2 incorporation to PRC2 **(Figure 3A)**. This was corroborated by *in vitro* stoichiometric assembly of a recombinant PRC2 core complex containing SUZ12, EED, RBBP4, and EZH2 -/+ SBD **(Supplementary Figure 3F,G).** Since EZH2 ΔSBD mediated ineffective H3K27me2/3 deposition *in vivo*, we next evaluated PRC2 recruitment to chromatin. ChIP-seq of EZH2 and SUZ12 in various mESC populations (EZH2 WT, KO and ΔSBD) yielded no difference in chromatin occupancy, other than the expected loss of EZH2 in <u>the</u> KO background **(Figure 3B,C)**. To assess any EZH2 requirement for the *in vivo* stability of PRC2 accessory subunits, we performed chromatin fractionation followed by immunoblotting MTF2, EPOP, AEBP2 and JARID2. As for PRC2 core subunits, the levels of chromatin-bound accessory proteins were similar in EZH2 WT, KO and ΔSBD mESCs **(Figure 3D)**. This was also observed by ChIP-seq of JARID2 and MTF2 in each EZH2 mESC background **(Supplemental Figure 3A)**. Together these data support the notion that SBD deletion does not impact PRC2 complex composition or recruitment to chromatin.

**Figure 3.**
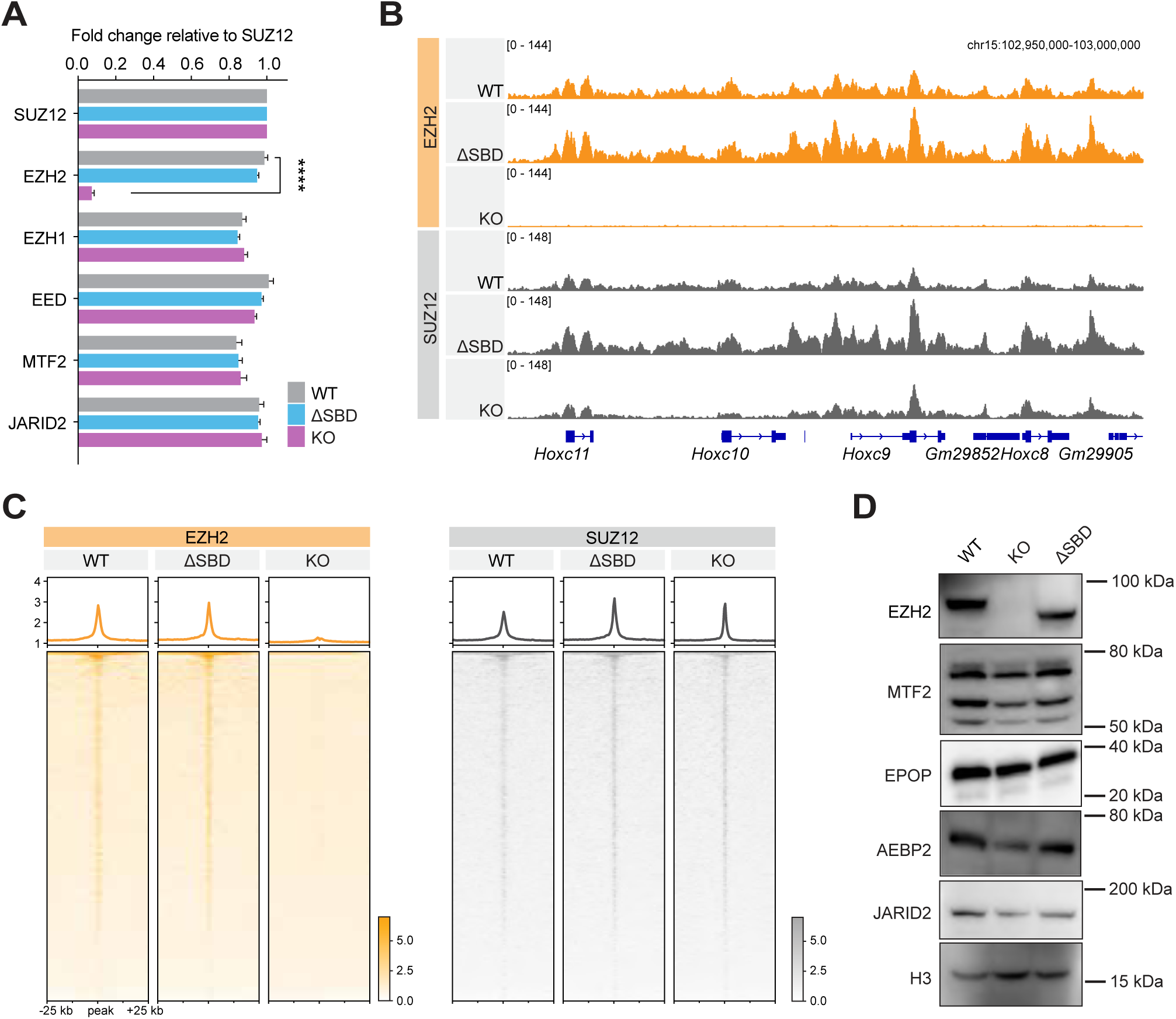
Deletion of EZH2 SBD does not compromise PRC2 complex integrity or recruitment to chromatin. **A.** Tandem MS analysis after SUZ12 co-IP whole nuclear extracts from EZH2 WT and EZH2-ΔSBD mESCs. The fold change of each subunit was calculated relative to SUZ12. Error bars = SEM. **** p-value = < 0.0001, the remaining comparisons were not significant by one-way ANOVA adjusted for multiple comparisons. **B.** Suz12 and EZH2 occupancy at *HoxC* locus by ChIP-seq. **C.** Heatmaps representing EZH2 (orange) and Suz12 (grey) ChIP-seq peaks centered on maximum peak signal -/+ 25-kb. **D.** Immunoblot of PRC2 accessory proteins after chromatin fractionation of mESC cells (EZH2 WT, KO or ΔSBD). H3 was used as a loading control.

To further establish the functional significance of the EZH2 SBD in regulating PRC2 activity, we performed *in vitro* histone methyltransferase assays using nuclear extracts from WT, KO and ΔSBD mESCs **(Supplementary Figure 3B,C)**. Here ΔSBD caused a dramatic reduction in PRC2 activity, similar to the effect seen with complete knockout of EZH2 methyltransferase **(Supplementary Figure 3B,C)**. This was corroborated using recombinant PRC2 WT and ΔSBD core complexes in a dCypher methylation assay of nucleosome substrates, where ΔSBD was inactive at baseline (**Supplementary Figure 3D**) and could not be allosterically activated by methylated JARID2 or H3 peptides (**Supplementary Figure 3E**).

Together, these results demonstrate that PRC2 recruitment to target genes is uncoupled from its enzymatic activity. Consequently, while the EZH2 SBD is dispensable for PRC2 complex assembly or recruitment to chromatin, it is essential for the formation of repressive H3K27me3 domains.

### EZH2 SBD is required for catalytic activity of the cancer-associated mutation EZH2 Y641N

Studies of PRC2 regulation in the context of oncogenic *EZH2* GOF mutations show that preventing allosteric activation can normalize H3K27me3 levels and suppress DLBCL proliferation ^5^. We therefore hypothesized that ΔSBD might similarly suppress the growth or tumorigenic effects of the *EZH2* Y641N GOF in mESCs and human lymphoma cell lines. To this end, we first ectopically expressed N-terminal FLAG tagged EZH2 Y641N or EZH2 ΔSBD-Y641N in mESCs expressing endogenous wild-type EZH2 (*EZH2* ^WT/WT^). By immunoblot of H3K27me2 and H3K27me3 bulk levels, ΔSBD prevented the increase of H3K27me3 driven by Y641N **(Supplementary Figure 5A)**. By quantitative ChIP-Rx, ectopic EZH2 Y641N led to a genome-wide increase in H3K27me3 and spreading of the mark beyond its wild-type boundaries **(Supplemental Figure 5B, C)**. Conversely, *EZH2* ΔSBD-Y641N maintained localization at wild-type levels **(Supplementary Figure 5B,C)**, indicating the SBD is needed to drive epigenomic changes caused by the EZH2 GOF mutant.

### EZH2 SBD is required to sustain proliferation of PRC2-addicted lymphoma cells

We next examined whether EZH2 SBD is required for the proliferation of cell lines derived from germinal center B-cell (GCB-type) lymphomas carrying EZH2 GOF mutations ^28^ **(Figure 4)**. To this end, we introduced an inducible knock-down system targeting endogenous EZH2 in a PRC2- dependent EZH2 Y641N line (Karpas-422), and a PRC2-independent EZH2 WT line (SU-DHL-8) **(Figure 4A)**, yielding the predicted H3K27me3 decrease in both lympoma backgrounds **(Supplementary Figure 4).** We then transformed each EZH2 knock-down line with ectopic rescue constructs (EZH2 WT, ΔSBD, Y641N or ΔSBD-Y641N) and performed *in vitro* differential growth analyses **(Figure 4)**. Here Karpas-422 cell proliferation was restored by Y641N but not ΔSBD-Y641N **(Figure 4B)**, while PRC2-independent SU-DHL-8 cells showed no response to ectopic expression of any EZH2 construct **(Figure 4C)**. To determine if loss of the EZH2 SBD would inhibit DLBCL proliferation *in vivo*, we transplanted engineered Karpas-422 cells to NSG mice and induced shRNA knockdown when tumors became palpable. Deletion of the SBD in the context of EZH2 Y641N led to tumor regression, similar to if the GOF allele was completely deleted **(Figure 4D)**. Such findings further emphasized that the SBD plays a critical role in EZH2 activity *in vivo*, including in the context of a disease-driving GOF mutation.

**Figure 4.**
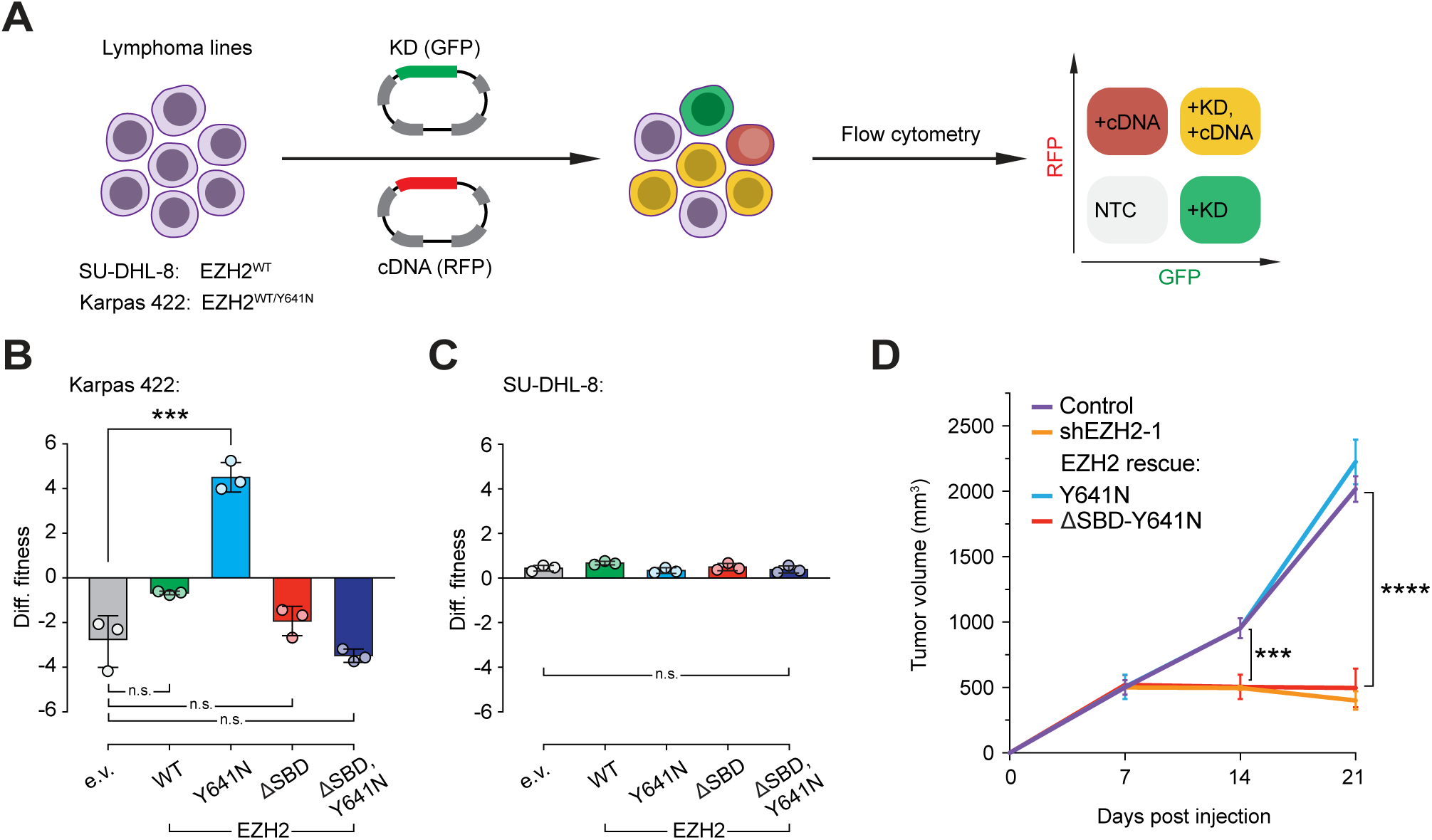
EZH2 SBD is required to support the growth of PRC2-dependent lymphomas. **A.** Outline of the experimental approach: wild type EZH2 (SU-DHL-8) and EZH2^+/Y641N^ gain of function mutant (Karpas-422) cells are transduced with GFP-labeled inducible knock-down construct and RFP-labeled rescue *EZH2* cDNA. Accumulation of GFP and RFP-labeled populations provides the fitness readout. **B.** Differential fitness after a 10-day competition assay in PRC2-dependent Karpas-422 cells. Error bars = mean ± SEM, p values calculated by Student’s *t* test. *** p-value = < 0.001, ns = not significant. **C.** Differential fitness after a 10-day competition assay in control SU-DHL-8 cells. Error bars = mean ± SEM, p values calculated by Student’s *t* test. *** p-value = < 0.001, ns = not significant. **D.** The impact of EZH2 ΔSBD on tumor growth was assessed in a Karpas-422 subcutaneous xenograft mouse model (n= eight flanks per cohort). Tumor volume is plotted as a function of time. Data is represented as the mean tumor volume per cohort and time point ± SD. P values were calculated by Two-way ANOVA with post hoc Bonferroni correction. *** p-value = < 0.001, **** p-value = < 0.0001

To assess if PRC2-mediated pathways drove the observed inhibition of proliferation we next examined the H3K27me3 landscape and transcriptomic profiles of various Karpas-422 cell lines. By CUT&RUN the control parental and [shEZH2; EZH2 Y641N transformed] lines showed similar H3K27me3 patterns, while the [shEZH2; ΔSBD-Y641N transformed] line exhibited genome-wide redistribution of H3K27me3 **(Figure 5A)**, with the PTM lost at known Polycomb target genes **(Figure 5B, C; Supplementary Figure 5)**. To further define the EZH2 SBD dependency of Karpas-422, we analyzed differentially expressed genes and H3K27me3 CUT&RUN data to the epigenomic landscape after pharmacologic inhibition of EZH2 with CPI-360 ^53^, identifying high similarity in the gene expression signature **(Figure 5D,E**; down- and upregulated genes overlapped with a normalized enrichment score of −2.3 and 1.76 respectively). Together, these data demonstrate the dependency of an EZH2 GOF lymphoma on an intact SBD function.

**Figure 5.**
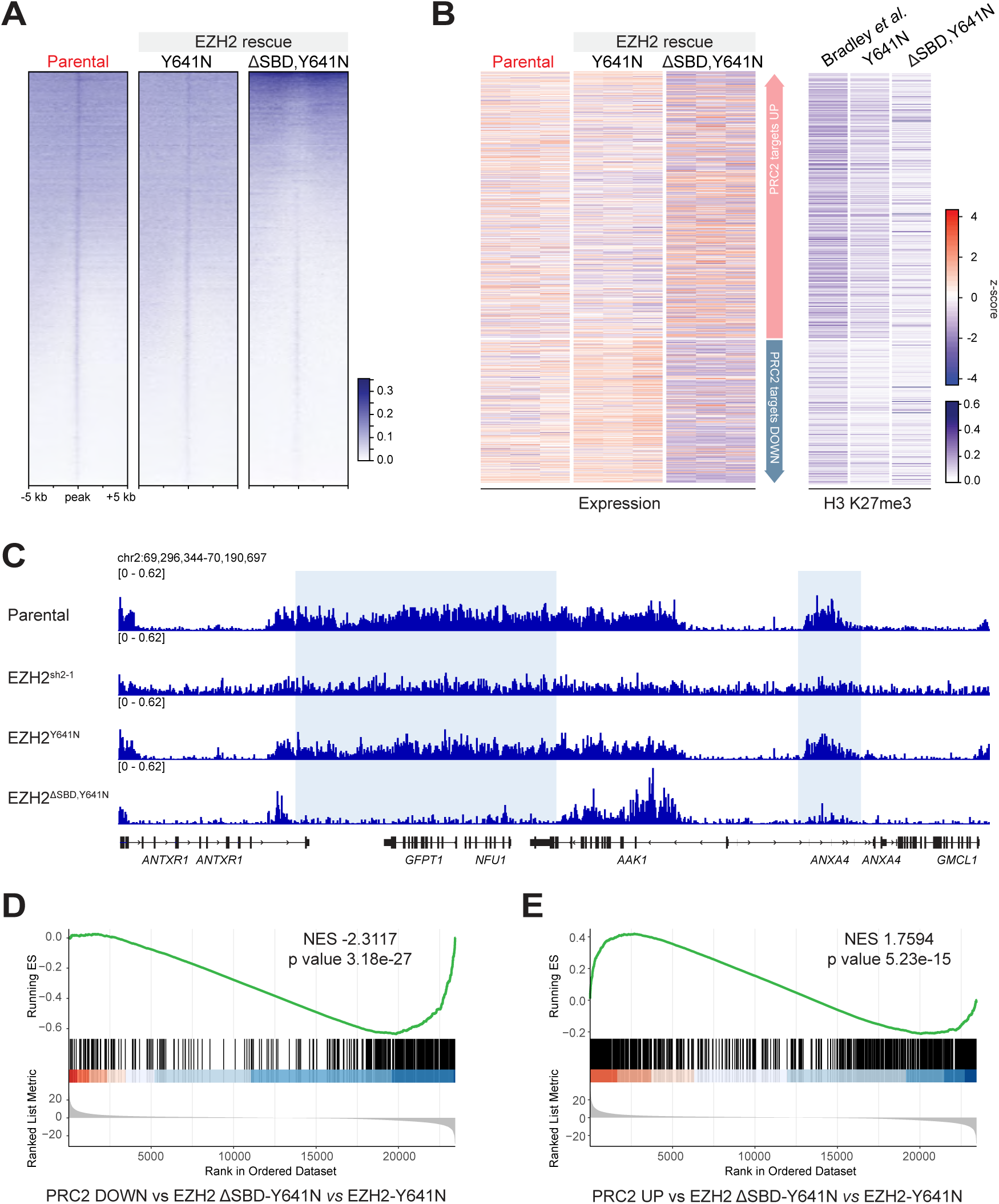
SBD is indispensable for the phenotype of EZH2 Y641 gain-of-function mutant. **A.** Heatmaps show H3K27me3 CUT&RUN signal (maximum peak signal -/+ 10kb) in parental Karpas-422 cells. **B.** Heatmaps show Z-scores of differentially expressed genes upon ectopic expression of EZH2-Y641N and EZH2-ΔSBD- Y641N (left), along with H3K27me3 signal (right) at Polycomb target genes from (Bradley *et al*. 2014). **C.** Genome browser representation of H3K27me3 CUT&RUN signal. Blue boxes highlight Polycomb targets in DLBCL. Refseq annotations are shown below. **D, E.** Gene Set Enrichment Analysis (GSEA) of genes down- (**D**) and up-regulated (**E**) after EZH2 inhibition in DLBCL cells, and their correlation with gene expression changes in cells rescued with EZH2-ΔSBD-Y641N.

## DISCUSSION

The PRC2 chromatin regulator plays a crucial role in developmental processes, primarily by forming and preserving cellular identity. Essential aspects of PRC2 function include controlled targeting to specific genomic regions, and managed histone methyltransferase (HTMase) activity at these locations. Here we show that the evolutionarily conserved EZH2 N-terminal SBD (SANT1-binding domain) is required to conserve the H3K27-methylation landscape.

Our data supports a model where the EZH2 SBD is not required for PRC2 recruitment to chromatin nucleation sites; but rather ΔSBD reduces PRC2 enzymatic activity, leading to diminished H3K27me3 domain formation **(Figure 6)**. The mouse EZH2 SBD is required to drive cell fate decisions during mESC differentiation, and enable chromatin spreading of both H3K27me2 and H3K27me3. However, the wild-type and ΔSBD PRC2 are similarly assembled and chromatin recruited, indicating the SBD is not required for complex stability or genomic localization. Of note, the few discrete loci that preserve H3K27 methylation in ΔSBD cells correspond to strong PRC2 nucleation sites enriched with CpG islands ^11^. Similar focal retention of H3K27me3 has been reported in H3K27M-driven malignancies ^22,54^, but to what degree SBD deletion recapitulates dominant PRC2 inhibition by “oncohistone” mutants remains to be determined.

**Figure 6.**
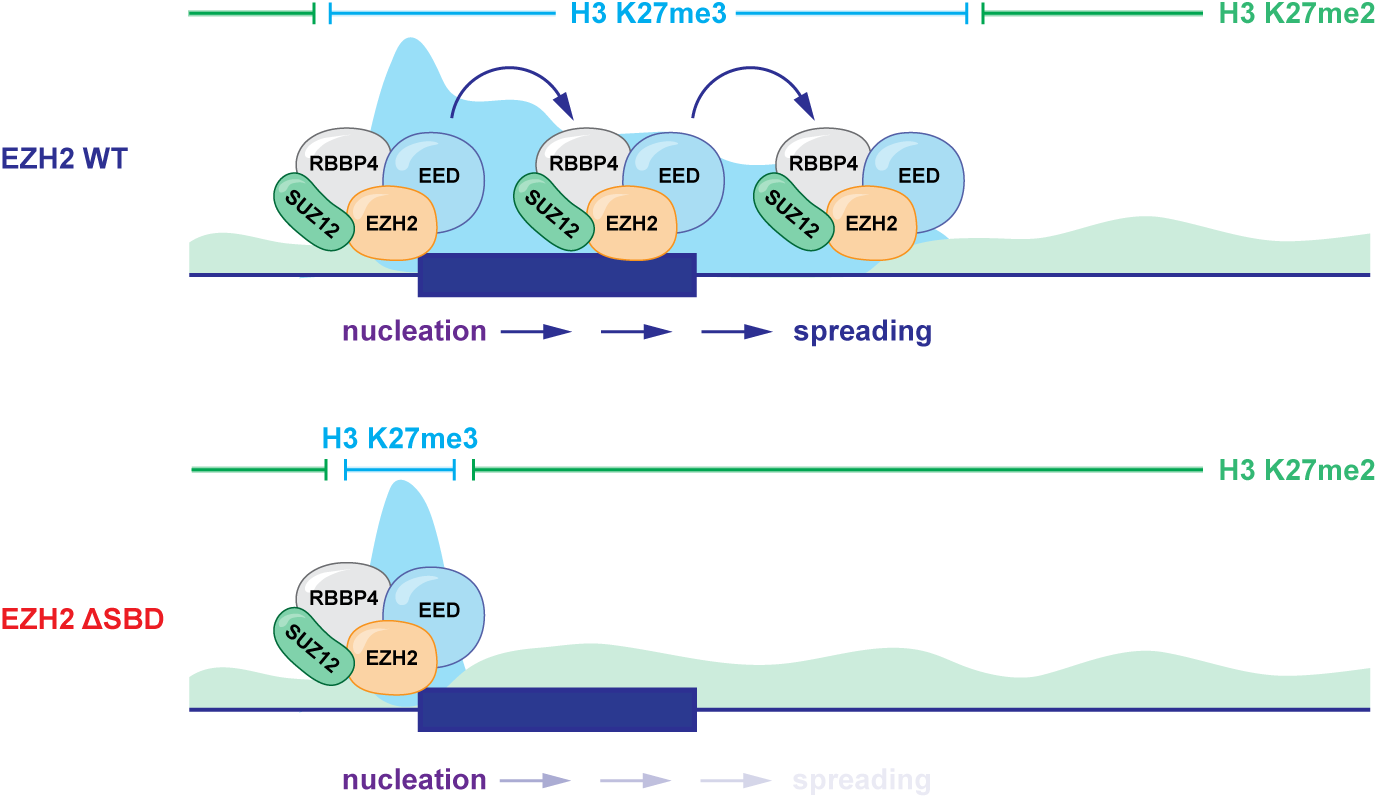
Model for inhibition of H3K27me3 spreading by EZH2-ΔSBD. EZH2 (WT) is recruited to PRC2 nucleation sites, where it catalyzes H3K27me3 and initiates PRC2 allosteric activation, leading to the formation of broad H3K27me3 domains. In contrast, EZH2-ΔSBD is similarly recruited to the PRC2 nucleation site but fails to catalyze sufficient H3K27me3 levels for allosteric activation. This results in PRC2 stalling at the nucleation site and the loss of broad chromatin domains.

We also investigated EZH2 SBD function in the context of tumorigenesis driven by the EZH2 GOF Y641N. PRC2 is frequently dysregulated in malignancy ^55^, with diffuse large B-cell lymphomas (DLBCLs) carrying recurrent EZH2 Y641N mutations that alter enzyme substrate specificity ^27,56^. Whereas wild-type EZH2 has an inherent preference to catalyze H3K27me1 and H3K27me2, Y641N promotes the conversion of H3K27me2 to H3K27me3. This deregulation of catalytic activity is central to oncogenesis in a subset of DLBCLs, highlighting the importance of controlling the rate-limiting step and balancing endogenous H3K27me3 levels in disease ^4–6,27^. Our work demonstrates that an intact EZH2 SBD is critical for Y641N GOF activity, where genetic inactivation of the SBD in GOF- addicted DLBCL cells both normalized H3K27me3 levels and suppressed cell proliferation. Further, this genetic manipulation had similar transcriptomic and epigenomic impact as a therapeutic intervention (CPI-360) targeting the EZH2 SET domain ^29^.

EZH2 posttranslational modifications may play an essential role in regulating its function, enzymatic activity and PRC2 complex stability ^57–60^. Curiously, phosphorylation of S21 within the EZH2 SBD has been shown to dissociate EZH2 from PRC2 and switch its function to a transcriptional coactivator ^60^, an observation corroborated by findings of a PRC2-independent role for EZH2 in transcriptional activation via Myc/p300^34^. While we have demonstrated an unexpected role for the spatially distant EZH2 SBD in regulating H3K27 methylation activity, our study did not discern any impact of ΔSBD on EZH2 complexation or PRC2 composition. In this vein, PRC2 activation through *trans-* autoactivation via dimerization was recently described ^57^, but further investigation will be required to determine if the SBD is implicated in this mode of enzymatic regulation. Finally, the discovery that SBD deletion can normalize EZH2 Y641N hyperactivity and impair DLBCL proliferation suggests that protein surfaces distant from the catalytic SET domain may provide additional therapeutic targeting strategies in PRC2/EZH2-addicted malignancy.

## Resource availability

### Lead contact

Further information and requests should be addressed to the lead contact, Agata Lemiesz Patriotis.

### Materials availability

The materials generated in this study are available upon the request to the lead contact.

### Data and code availability

- All sequencing data created in this study will be uploaded to the Gene Expression Omnibus (GEO; https://www.ncbi.nlm.nih.gov/geo/) and are available under accession code GEO: GSE286456, GSE286457, GSE286459, GSE286460, GSE286462.
- This study does not report the original code. Publicly available software is listed in the methods section.
- Any additional information required to re-analyze results reported in this manuscript is available upon request to the <u>lead contact</u>.

## Acknowledgments

We thank colleagues for the generous gift of reagents (see **Experimental Procedures**) and members of the Allis, Garcia, and Muir laboratories, Shixin Liu and Richard Lifton for helpful discussions and support. A.L.P. was supported by the Women & Science Fellowship at the Rockefeller University, Anderson Cancer Center Fellowship for Cancer Research at the Rockefeller University, and an F99/K00 Predoctoral to Postdoctoral Fellowship from the National Cancer Institute (F99CA253687). Y.S.F was supported by a Damon Runyon-Sohn Pediatric Cancer Fellowship (DRSG 21-17) and a National Institutes of Health (NIH) / NIGMS K99/R00 Award (1K99GM140265). A.A.S. is supported by R01 CA234561, American Cancer Society IRG-21-147-01-IRG, CPRIT IIRA (RP240068) and HIHR (RP240446), and institutional funds from the University of Texas at San Antonio. *EpiCypher* was supported by NIH grants R44GM117683 and R44GM116584. C.D.A. was supported by NIH grants (P01CA196539 and 5R01CA204639), The Leukemia & Lymphoma Society (LLS-SCOR 7006-13), and The Rockefeller University and St. Jude Children’s Research Hospital Collaborative on Chromatin Regulation in Pediatric Cancer. We acknowledge help and support from the Bio-Imaging (RRID:SCR_017791), Genomics (RRID:SCR_020986), Proteomics (RRID:SCR_017787), Flow Cytometry (RRID:SCR_017694), Bioinformatics, and Comparative Bioscience Resource Centers at The Rockefeller University.

## Competing interests

*EpiCypher* is a commercial developer and supplier of nucleosomes and the *dCypher* platform used in this study. L.K. M.R.M. and M.-C.K. are employed by (and own shares in) *EpiCypher*, with M.-C.K. also a board member of same. The authors declare no other competing interests.

## Author contributions

A.L.P., A.A.S. and C.D.A. conceptualized this project. A.L.P. wrote the manuscript with input from all authors. A.L.P., A.A.S., L.K., A.D., P.L., D.M., Y.S.F., L.K, M.R.W. and D.B. generated reagents, conducted experiments, and/or analyzed and interpreted data. M.-C.K., T.S.C., B.A.G., and C.D.A. supervised the study.

## Experimental procedures

### Gene and protein nomenclature

EZH2 protein and its N-terminal region are highly conserved across species. We therefore chose to refer to both mouse and human EZH2 protein using uppercase notation to simplify readability. Where applicable, the human *EZH2* gene is denoted in all uppercase italics, while for the mouse *Ezh2* gene only the first letter is capitalized. Other genes and proteins are denoted according to conventional VGNC nomenclature.

### Plasmid generation and lentivirus production for cell culture

Single guide (sgRNAs) directed against mouse *Ezh2* were cloned into PX458 (Addgene 48138; a gift from F. Zhang). Mouse *Ezh2* cDNA sequences from Horizon Dharmacon were cloned into PiggyBac (pCAGGS-IRES-Neo; a gift from H. Niwa, Institute of Molecular Embryology and Genetics, Kumamoto) and pCDH-EF1-MCS-IRES-GFP (System Biosciences, CD531A-2). The amino-terminal region of mouse EZH2 containing the SBD domain (1-23 aa) were replaced with amino-terminal sequences of EZH2 orthologs as described in Supplemental Figure 1. All cloning including domain-deletion, domain-swap, and patient-associated oncohistone mutations was performed using Gibson assembly (NEB). To produce lentivirus, 293T cells were transfected with the lentiviral vector and helper plasmids (psPAX2, pVSVG). Supernatant containing lentivirus was collected and filtered 48 hours later for transduction.

### Cell culture, CRISPR-Cas9 gene editing and generation of stable cell lines

V6.5 mouse ES cells (C57BL/6 × 129S4/SvJae F1) were maintained on gelatin-coated plates in KnockOut DMEM (Gibco) supplemented with 15% ES-cell-qualified FBS (Gemini), 0.1 mM 2- mercapoethanol, 2mM L-glutamine (Life Technologies) and LIF. 293T cells (ATCC) were cultured in DMEM supplemented with 4.5 g/l glucose (Corning), 10% FBS (Atlanta Biologicals), 2 mM L-glutamine (Life Technologies) and 1:100 penicilin/streptomycin (Corning). *Drosophila* S2 cells were cultured in Schneider’s Drosophila medium (Gibco) containing 10% heat-inactivated FBS; (Atlanta Biologicals, S11150, lot C19031). Karpas-422 (Sigma) and SU-DHL-8 (ATCC) were cultured in suspension in RPMI-1640 (Invitrogen) supplemented with 20% FBS (Sigma). All cell lines were regularly confirmed as negative for mycoplasma contamination. To generate knockout lines, mouse ES cells were transfected with sgRNA-containing PX458 using Xfect mouse ES cell transfection reagent (Takara) and incubated for 48 h. Single GFP^+^ cells were then sorted into 96-well plates.

Clones were expanded, screened for global reduction of H3K27me3 by immunoblot and individually verified by Sanger sequencing of the target loci. To generate transgenic mouse ES cell lines, around 5 × 10^6^ cells were electroporated with PiggyBac expression vectors plus transposase (pBase) in a 3:1 ratio using the Amaxa ESC Nucleofector kit (VPH-1001, program A-023, Lonza). Cells were plated on gelatin-coated plates and grown under G418 (500 μg/mL) selection 48 h after electroporation for at least two passages before being collected for immunoblot or ChIP–seq analysis. To generate transduced Karpas-422 and SU-DHL-8 lines expressing EZH2 mutants, 500 k cells were plated per well of a 12-well non-TC treated plate in 2 mL media, lentivirus and polybrene (10μg/mL). The plates were centrifuged at 1300 g for 90 min at 37°C. Cells were cultured in non-TC treated 6-well plates, puromycin (Invivogen; 1μg/mL) added 48 hrs after transduction, and selected for at least 10 days, with appropriate negative controls.

### Histone acid extraction, histone derivatization and analysis of post-translational modifications by nano-LC–MS

Mouse ES cells were lysed in nuclear isolation buffer (15 mM Tris pH 7.5, 60 mM KCl, 15 mM NaCl, 5 mM MgCl_2_, 1 mM CaCl_2_, 250 mM sucrose, 10 mM sodium butyrate, 1 mM DTT, 500 µM AEBSF and 5 nM microcystin) containing 0.3% NP-40 alternative on ice for 5 min. Nuclei were pelleted and resuspended in 0.2 M H_2_SO_4_, followed by 1.5 h rotation at 4 °C. After centrifugation, supernatants were collected and proteins precipitated in 33% TCA overnight on ice, washed with acetone, and resuspended in deionized water. Acid-extracted histones (5-10 μg) were resuspended in 100 mM ammonium bicarbonate (pH 8), derivatized using propionic anhydride and digested with trypsin as described previously ^61^. After a second round of propionylation, the resulting histone peptides were desalted using C18 Stage Tips, dried using a centrifugal evaporator, and reconstituted using 0.1% formic acid in preparation for liquid chromatography–mass spectrometry (LC–MS). Nanoflow liquid chromatography was performed using an Easy nLC 1000 (Thermo Fisher Scientific) equipped with a 75 µm × 20-cm column packed in-house using Reprosil-Pur C18-AQ (3 µm; Dr. Maisch). Buffer A was 0.1% formic acid and Buffer B was 0.1% formic acid in 80% acetonitrile. Peptides were resolved using a two-step linear gradient from 5% B to 33% B over 45 min, then from 33% B to 90% B over 10 min at a flow rate of 30 nL/min. The HPLC was coupled online to an Orbitrap Elite mass spectrometer operating in the positive mode using a Nanospray Flex Ion Source (Thermo Fisher Scientific) at 2.3 kV. Two full mass spectrometry scans (m/z 300–1,100) were acquired in the Orbitrap mass analyser with a resolution of 120,000 (at 200 m/z) every 8 data-independent acquisition tandem mass spectrometry (MS/MS) events, using isolation windows of 50 m/z each (for example, 300–350, 350–400…650–700). MS/MS spectra were acquired in the ion trap operating in normal mode. Fragmentation was performed using collision-induced dissociation in the ion trap mass analyzer with a normalized collision energy of 35. The automatic gain control target and maximum injection time were respectively 5 × 10^5^ and 50 ms for the full mass spectrometry scan, and 3 × 10^4^ and 50 ms for the MS/MS scan. Raw files were analyzed using EpiProfile 2.032. The area for each modification state of a peptide was normalized against the total signal for that peptide to give relative abundance of the histone modification.

### Chromatin fractionation

Mouse ES cells were washed with PBS and lysed on ice for 8 min in Buffer A (10 mM HEPES, 10 mM KCl, 1.5 mM MgCl2, 0.34 M sucrose, 10% glycerol, 0.5 mM PMSF and 0.1% Triton X-100). Centrifugation was performed (1,300g at 4 °C for 5 min) and the supernatant collected (cytosolic fraction). The nuclei pellet was further lysed on ice for 30 min in Buffer B (3 mM EDTA, 0.2 mM EGTA and 0.2 mM PMSF) and centrifugation performed (1,700g at 4 °C for 5 min) to obtain the supernatant (nuclear soluble fraction). Cytosolic and nuclear soluble fractions were combined to make the soluble fraction. The insoluble pellet was lysed in SDS sample loading buffer, boiled, and sonicated to yield the chromatin fraction. The protein concentration of each fraction was measured, and equal amounts analyzed by immunoblot.

### Immunoblotting

Fractionated or whole cell lysates were resolved by SDS-PAGE, transferred to a nitrocellulose or PVDF membrane, blocked in 5% non-fat milk in PBS plus 0.5% Tween-20, probed with primary antibodies, and detected with horseradish peroxidase (HRP)- conjugated anti-rabbit or anti-mouse secondary antibodies (GE Healthcare). Primary antibodies were: anti-H3K27me2 (CST, 9728), anti-H3K27me3 (CST, 9733), anti-SUZ12 (CST, 3737), anti-EZH2 (CST, 3147), anti-MTF2 (Proteintech, 16208-I-AP), anti-AEBP2 (CST, 14129), anti-JARID2 (CST, 13594), anti-EED (Millipore, 09-774), anti-H2B (CST, 12364), anti-β-Tubulin (CST, 2128), anti-β-actin (Abcam, ab8224), anti-H3 (Abcam, ab1791) and anti-HA (Biolegend, 901501).

### PRC2 enzymatic assays

PRC2 methyltransferase activity assays were performed using AlphaScreen technology (Revvity). To compare the enzymatic activity of Ezh2 WT and ΔSBD complexes, each were serially titrated in reaction buffer (20 mM Tris pH 7.5, 0.01% BSA, 0.01% Tween-20, 1 mM DTT). Enzyme (2.5 uL of dilution), nucleosome substrate (2.5 μL of 10 nM biotinylated unmodified; EpiCypher 16-0006), allosteric activator (2.5 μL of 6.25 μM JARID2_[109-123]_K116me3 peptide) and S-Adenosyl Methionine (SAM) co-factor (2.5 μL of 400 μM) were combined to 10 uL in a 384-well plate (Revvity, 60053595) and reactions incubated for two hours at 23 °C. H3K27me1 product was detected by addition of 10μL anti-H3K27me1 (RevMab Clone#27; diluted 1:20,480 in antibody buffer (625 mM NaCl, 20 mM Tris pH 7.5, 0.01% BSA, 0.01% Tween-20, 1 mM DTT). After antibody addition and incubation at 23 °C for 30 min, a 5 μL mix of protein A acceptor (Revvity AL101C; 12.5 μg/mL) and streptavidin donor (Revvity 6760002; 50 μg/mL) beads in reaction buffer was added to each well, and further incubated for one hour at 23 °C (protecting from light from this step). Alpha signal was measured on a PerkinElmer 2104 EnVision (680 nm laser excitation, 570 nm emission filter ± 50 nm bandwidth), and data analyzed in GraphPad Prism 10 using four parameter logistic non-linear regression. Each reaction was performed in duplicate.

Experiments to examine allosteric activation of Ezh2 WT and ΔSBD complexes (both 12.5 nM) was compared using a serial titration of JARID2_[109-123]_K116me3 and H3_[23-34]_K27me3 peptides.

### Chromatin Immunoprecipitation (ChIP)

Cross-linking ChIP in mESCs was performed as described ^62^ using ∼2 × 10^7^ cells per immunoprecipitation. Before fixation, medium was aspirated, cells were collected by TrypLE Express Enzyme (Gibco), and cells washed once with PBS. Cells were cross-linked (1% paraformaldehyde for 5 min at room temperature (RT°)) with gentle shaking. Glycine was added to quench (final concentration 125 mM, incubated for 5 min at RT°), then cells washed once with cold PBS and pelleted. To obtain a soluble chromatin extract, cells were resuspended in 1 ml LB1 (50 mM HEPES, 140 mM NaCl, 1 mM EDTA, 10% glycerol, 0.5% NP-40, 0.25% Triton X-100 and 1× Complete protease inhibitor) and incubated rotating at 4 °C for 10 min. Samples were centrifuged, resuspended in 1 ml LB2 (10 mM Tris-HCl pH 8.0, 200 mM NaCl, 1 mM EDTA, 0.5 mM EGTA and 1× Compete protease inhibitor), and incubated rotating at 4 °C for 10 min. Finally, samples were centrifuged, resuspended in 1 ml LB3 (10 mM Tris-HCl pH 8.0, 100 mM NaCl, 1 mM EDTA, 0.5 mM EGTA, 0.1% Na deoxycholate, 0.5% N-lauroylsarcosine, 1% Triton X-100 and 1× Complete protease inhibitor) and homogenized by passing two times through a 27-gauge needle. Chromatin extracts were sonicated for 14 min using a Covaris E220 focused ultrasonicator at peak power 140, duty factor 5, and cycles/burst 200. For ChIP of histone post-translational modifications, after centrifugation, samples were spiked with soluble chromatin from *Drosophila* S2 cells to comprise 2–5% of total chromatin in the lysate. The lysates were 75 μl protein A Dynabeads (Invitrogen) bound to anti-H2AK27me2 (CST, 9728), anti-H3K27me3 (CST, 9733), anti-SUZ12 (CST, 3737) or anti-EZH2 (CST, 5246) [all at 1:50 dilution], and incubated overnight at 4 °C, with 5% kept as input DNA. Magnetic beads were sequentially washed with low-salt buffer (150 mM NaCl; 0.1% SDS; 1% Triton X-100; 1 mM EDTA and 50 mM Tris-HCl), high salt buffer (500 mM NaCl; 0.1% SDS; 1% Triton X- 100; 1 mM EDTA and 50 mM Tris-HCl), LiCl buffer (150 mM LiCl; 0.5% Na deoxycholate; 0.1% SDS; 1% Nonidet P-40; 1 mM EDTA and 50 mM Tris-HCl) and TE buffer (1 mM EDTA and 10 mM Tris-HCl). Beads were resuspended in elution buffer (1% SDS, 50 mM Tris-HCl pH 8.0, 10 mM EDTA and 200 mM NaCl) and incubated for 30 min at 65 °C. After centrifugation, the eluate was reverse cross-linked overnight at 65 °C. The eluate was then sequentially treated with RNase A for 1 h at 37 °C and Proteinase K (Roche) for 1 h at 55 °C, and DNA recovered using a Qiagen PCR purification kit.

### CUT&RUN

CUT&RUN was performed with minor modifications of the CUTANA (*EpiCypher*) protocol ^63–65^. Briefly, cells were fixed with 0.1% formaldehyde in PBS for 1 minute, followed by quenching with 125 mM glycine. Cells were washed with cold Wash Buffer (20 mM HEPES-KOH, pH 7.9, 150 mM NaCl, 0.5 mM Spermidine, protease inhibitor cocktail) and then bound to concavalin A beads for 10 minutes at room temperature. Supernatant was removed and beads were resuspended in Antibody Buffer (20 mM HEPES-KOH, pH 7.9, 150 mM NaCl, 0.5 mM Spermidine, 0.01% Digitonin, 2 mM EDTA, protease inhibitor cocktail) along with primary antibody. Samples were incubated overnight in the cold room, rotating. Beads were then resuspended in cold Digitonin Buffer 20 mM HEPES-KOH, pH 7.9, 150 mM NaCl, 0.5 mM Spermidine, 0.01% Digitonin, protease inhibitor cocktail), washed twice, then incubated with CUTANA pAG-MNase (Epicypher) for 10 minutes at RT. Beads were then washed twice with cold Digitonin buffer, resuspended in Digitonin Buffer with 50 mM CaCl_2_, and samples incubated for two hours with rotation at 4 °C. MNase was quenched by addition of STOP Buffer (340 mM NaCl, 20 mM EDTA, 4 mM EGTA, 50 mg/ml RNase, 50 mg/ml Glycogen) and incubation for 10 mins at 37 °C. 0.1% SDS and Proteinase K was added to the supernatant and incubated for one hour at 55 °C followed by two hours at 65 °C. DNA was purified using Phenol-Chloroform Isoamyl Alcohol mix (Millipore) and PhaseLock tubes (5PRIME), and quantified using a Qubit 4 Fluorometer (Thermo) prior to proceeding with library preparation.

### RNA isolation

Total RNA was isolated from cultured cells using RNeasy Mini Kit (Qiagen) with on-column DNA digestion. RNA samples were analyzed on the Bioanalyzer RNA 6000 Pico (Agilent) prior to library preparation.

### Library preparations & sequencing

RNA libraries were converted to cDNA using NEBNext Poly(A) mRNA Magnetic Isolation Module (NEB) and NEBNext Ultra II RNA Library Prep Kit for Illumina (NEB) according to the manufacturer’s instructions. ChIP and CUT&RUN libraries were prepared using NEBNext Ultra II DNA Library Prep Kit for Illumina (NEB) according to the manufacturer’s instructions. All libraries were analyzed by TapeStation prior to sequencing. Single-end sequencing was performed on the Illumina NextSeq 500 sequencer.

### Flow Cytometric Analyses

Population analyses were done by collecting lymphoma cells after treatment with Doxycycline and staining with conjugated primary antibodies (Pacific Blue anti-H3 (CST, 12167) or Alexa Fluor 647 anti-H3K27me3 (CST, 5499)) or analyzing for tRFP or eGFP. Stained samples were analyzed on an LSRFortessa (BD Biosciences) flow cytometer and data analysis performed using FlowJo (BD Biosciences). Intracellular antigen detection was done using the Foxp3/Transcription Factor Staining Buffer Set (eBioscience) per manufacturer’s guidelines.

### Growth Competition Assays

SU-DHL-8 and Karpas-422 cells expressing shRNA targeting EZH2 were virally transduced (in triplicate) with the designated rescue constructs (pCDH-cDNA-tRFP) in 12-well plates at 30% to 40% infection rate. Cells were monitored by flow cytometry over time using an LSRFortessa (BD Biosciences), and relative growth of rescue cDNA-containing cells assessed. Flow cytometry data were analyzed with FlowJo software (BD Biosciences). The percentage of single-positive (tRFP^+^ or eGFP^+^) or double-positive (tRFP^+^/eGFP^+^) cells was normalized to their respective “T_0_” time-point values. Normalized values were log_2_-transformed and relative cell proliferation calculated as *log_2_(normalized DP) − log_2_(normalized SP)*

### Cellular differentiation assays

Embryoid body differentiation was performed by plating 100 k mESCs per well of a six-well plate and growing for seven days in mES cell media lacking Leukemia Inhibitory Factor. Cells were washed with PBS and stained with Alkaline Phosphatase Assay Kit (Sigma Aldrich, MAK447).

### Statistical Analysis

All data are reported as the mean of experimental replicates with error bars designating either standard deviations or standard error of the mean as denoted in figure legends. Measurements were taken from distinct samples in all experiments. Statistical tests, the number of experimental replicates (n), and significance thresholds are described in figure legends.

### RNA-seq analysis

Transcript abundance was determined from FASTQ files using Salmon (v0.8.1) and the GENCODE reference transcript sequences ^66^. Transcript counts were imported into R with the tximport R Bioconductor package (v1.8.0), and differential gene expression performed with the DESeq2 R Bioconductor package (v1.20.0) ^67,68^. Normalized counts were retrieved from the DESeq2 results and z-scores for the indicated gene sets visualized with heatmaps generated using the ComplexHeatmap R Biconductor package (v 2.8.0) ^69^. Genes in heatmaps were annotated with ChIPseq signal by quantifying signal in regions starting at the transcription start site (TSS) and extending 5 kb downstream. Signal was quantified and then summarized as the mean signal in 50bp bins across these regions using the profileplyr R Bioconductor package (v1.8.1) ^70^). For gene set enrichment analysis (GSEA), the Wald statistic from the DESeq2 analysis was used to generate the ranked list of genes. The clusterProfiler R Bioconductor package (v4.0.5) was then used to perform GSEA to determine the enrichment of published up- and down-regulated polycomb genes within this ranked list ^71,72^ The enrichplot R Bioconductor package (v1.12.3) was used to visualize the GSEA result ^73^.

### ChIP-seq analysis

ChIP-seq reads were aligned using the Rsubread R Bioconductor package (v1.30.6) and predicted fragment lengths calculated by the ChIPQC R Bioconductor package (v1.16.2) ^74,75^. Normalized, fragment-extended signal bigWigs were created using the rtracklayer R Bioconductor package (v1.40.6), and peaks called using MACS2 (v2.1.1) ^76,77^. To determine the overlap of CpGs with H3K27me3 peaks, the loci of CpGs in the mouse genome (mm10) were obtained from the UCSC table browser (specifically the cpgIslandExt table from https://genome.ucsc.edu), and a bigWig file representing CpG coverage was generated with the rtracklayer R Bioconductor package. This bigwig file was then used to quantify CpG overlap of 20bp bins +/- 5 kb from the center of H3K27me3 peak regions using the profileplyr R Bioconductor package. The mean proportion of base pairs in each bin that overlap CpGs was then plotted using the generateProfilePlot function from profileplyr.

### Endogenous Co-Immunoprecipitations

Mouse embryonic stem cells were resuspended in Buffer C (420 mM NaCl, 20 mM HEPES pH 7.9, 0.2 mM EDTA, 1.5 mM MgCl_2_, 20% glycerol, 2 μg/mL Aprotonin, 1 μg/mL Leupeptin, 1 mM PMSF), sonicated 3x 15 seconds and dounced 20 times with a tight pestle. Lysates were incubated at 4°C for 20 min with rotation and clarified by centrifugation (20,817g) at 4°C for 20 min. Lysates were dialysed for five hours at 4°C against 50 volumes of Buffer C100 (125 mM KCl, 20 mM HEPES pH 7.9, 0.2 mM EDTA, 1.5 mM MgCl_2_, 20% glycerol). Lysates were again clarified by centrifugation (20,817g) at 4°C for 20 min. 5 μg SUZ12 antibody (Cell Signaling Technologies, 3737) was coupled to 20 μL magnetic Protein A beads (Invitrogen) in 1 mL PBS (0.1% Tween-20) overnight at 4°C with rotation. Beads were collected on the magnet at RT° and washed twice in 1 mL 0.2 M Sodium Borate pH 9.0. Beads were washed once in Buffer C100 and blocked for 60 minutes at 4°C, rotating in Buffer C100 (with 0.1 mg/mL Insulin (Sigma) and 0.2 mg/mL Chicken egg albumin (Sigma)). After the final wash, beads were resuspended in 100 μL SDS-PAGE sample buffer and immunoprecipitated material eluted by boiling for 5 min. Beads were vortexed, centrifuged at 20,817g for 5 minutes, and supernatant retained. For LC-MS/MS analysis, beads were flash-frozen in liquid nitrogen.

### Animals

NOD.Cg-Prkdcscid Il2rgtm1Wjl/SzJ 5-week-old female mice were purchased from The Jackson Laboratory. Each mouse (four per group) was inoculated in both flanks with the Karpas-422 tumor cells (5 x 10^6^) in 0.2ml of PBS with Matrigel (1:1) for tumor growth. Treatment with Doxycyline 625 irradiated diet (Envigo TD.01306) started seven days post-inoculation when average tumor size reached ∼ 100-200 mm^3^. The chow was replaced every two days for 14 days of treatement. Tumor size was measured three times a week using a caliper, and tumor volume (V, in mm^3^) expressed using V = 0.5a × b2, where a and b were respectively the long and short diameters. Animal care and use followed NIH guidelines and was approved by the Institutional Animal Care and Use Committee (IACUC) at The Rockefeller University.

