## Supplemental Figures for "The conserved N-terminal SANT1-binding domain (SBD) of EZH2 Regulates PRC2 Activity"

**Supplementary Figure 1. Ezh2 SBD is required for Polycomb regulation in mESCs. A.** Ezh2 SBD is highly conserved between EZH2 orthologs. Block height depicts cross-species SBD residue conservation. **B.** Immunoblot of mESCs lacking EZH2 (EZH2 KO), or individually complemented by EZH2-ΔSBD or EZH2-SBD orthologs from soft-shell turtle (*P. sinensis, Psi*), fruitfly *(D. melanogaster*, *Dme*) or *nematode (C. elegans*, *Cel*). Of note, the turtle (*Psi)* and fruitfly (*Dme)* alleles rescued H3K27me2 and H3K27me3 levels, while nematode (*Cel*) did not. β-actin and H3 were used as loading controls. **C.** Alkaline phosphatase (AP) staining of mESCs of indicated genotypes (where high AP activity is correlated to the inability of EZH2 to methylate H3K27).

**Supplementary Figure 2. EZH2 SBD is required for H3K27me3 broad domain formation. A.** Heatmaps show H3K27me3 in parental (EZH2 WT), EZH2 SET KO and EZH2-ΔSBD mESCs, and a published EED-rescue experiment[11] re-expression of EED at 12 and 36 hrs, EED Y365A, and EED WT. Plots are centered on maximum peak signal -/+ 10kb in WT mESCs. H3K27me3 peaks were sorted in descending order by signal intensity in EZH2-ΔSBD mESCs. Strong nucleation sites are marked in red, weak nucleation sites in blue, proximal spreading sites in yellow, and distal spreading sites in green. **B.** Nucleation sites preserve low levels of H3K27me3 in EZH2-ΔSBD mESCs. Representative genome browser views of EZH2 (orange), H3K27me3 (navy) and H3K27me2 (green) ChIP-Rx experiments in mESCs of three indicated genotypes. RefSeq annotations are below. Red highlight indicates nucleation site within the *Emx1* gene that preserved the H3K27me3 signal despite EZH2-KO or expression of EZH2-ΔSBD; residual H3K27me2 and H3K27me3 signals in Ezh2 KO cells reflect Ezh1 activity. **C.** Density plot of H3K27me3 signal overlapping with CpG islands in parental (EZH2 WT, black) and EZH2-ΔSBD (red) mESCs. Peaks were binned by 1 kb and the proportion of overlap with annotated CpG islands plotted. **D.** Heatmaps of H3K27me3 (blue) and H3K27me2 (green) ChIP-Rx signal, centered on the H3K27me3 maximum peak signal -/+ 10kb in WT (parental) mESCs. **E.** Boxplots of H3K27me3 ChIP-Rx signal genome-wide (top) and within peak regions mapped in parental (EZH2 WT) mESCs (bottom); p values are calculated by Student's *t*-test**.** ****, p-value = < 0.0001. **F.** Heatmap of Spearman's rank correlation coefficient values for each pairwise combination of ChIP-Rx experiments.

**Supplemental Figure 3. EZH2 SBD is not required for PRC2 complex assembly or the chromatin recruitment of MTF2 and JARID2 accessory subunits but is necessary for the enzymatic activity of PRC2 complex. A.** Heatmaps representing JARID2 (green) and MTF2 (orange) ChIP-seq peaks centered on maximum peak signal value -/+ 5kb. **B.** Histone methylation activity quantified by colorimetric readout. Error bars = SEM. *** p-value = < 0.001, by unpaired two-tailed t test. **C.** Varying quantities of nuclear extract were assayed for histone methyltransferase activity. Error bars = Mean + SD. Two-way ANOVA calculated P values with post hoc Bonferroni correction. *** p-value = < 0.001, **** p-value = < 0.0001. **D.** Recombinant PRC core complexes containing EZH2 WT and EZH2-ΔSBD were assayed by AlphaScreen for *in vitro* methylation of nucleosome substrates to H3K27me1 state in the presence of $JARID2_{[109-123]}$K116me3 allosteric activator peptide +/- SAM cofactor. **E.** Impact of titrating allosteric activating peptides $JARID2_{[109-123]}$K116me3 or $H3_{[23-34]}$K27me3 on methylation activity of EZH2 WT and EZH2-ΔSBD PRC2. **F.** Immunoblot of epitope tagged subunits of recombinant PRC2-EZH2-ΔSBD. **G.** SDS-PAGE / Coomassie stain demonstrates stoichiometry of recombinant PRC2-EZH2-ΔSBD.

**Supplementary Figure 4. Development of shEZH2 knock-down system to investigate the impact of SBD deletion on proliferation and tumorigenesis in B-cell lymphoma. A.** Experimental approach to generate inducible shEZH2 knock-down. Three different shRNAs were tested and shEZH2-1 chosen as the most effective (not shown). **B.** Flow cytometry analysis in SU-DHL-8 and KARPAS-422 cells expressing inducible shEZH2-1 ($GFP^{+}$). Note that nearly all SU-DHL-8 cells are GFP+ upon shEZH2 induction, while Karpas-422 cells maintain a large proportion of GFP- cells, suggesting strong dependency on EZH2. **C & D.** H3K27me3 loss after EZH2 knockdown in lymphoma cells. Intracellular staining coupled with flow cytometry for H3 **(C)** and H3K27me3 **(D)**. Error bars = SEM, p values are calculated by Student's *t*-test. ** p-value = < 0.01, * p-value = < 0.05, ns = not significant.

**Supplementary Figure 5. The EZH2 SBD is required for gain-of-function activity of EZH2-Y641N. A.** Immunoblot of mESCs expressing ectopic N-terminally tagged EZH2-Y641N or EZH-ΔSBD-Y641N (* truncated protein visible as a lower MW band). β-actin and H3 are loading controls. **B.** Representative genome browser view of H3K27me3 (navy) and H3K27me2 (green) ChIP-Rx signal. Refseq annotations are below. **C.** Heatmaps of H3K27me3 (navy) and H3K27me2 (green) ChIP-Rx signal (centered on peak maximum -/+ 5kb) in EZH2 WT mESCs. **D.** Box plot representing H3K27me3 ChIP-Rx levels in Karpas-422 cells within genes with altered expression upon EZH2 inhibitor treatment [53].

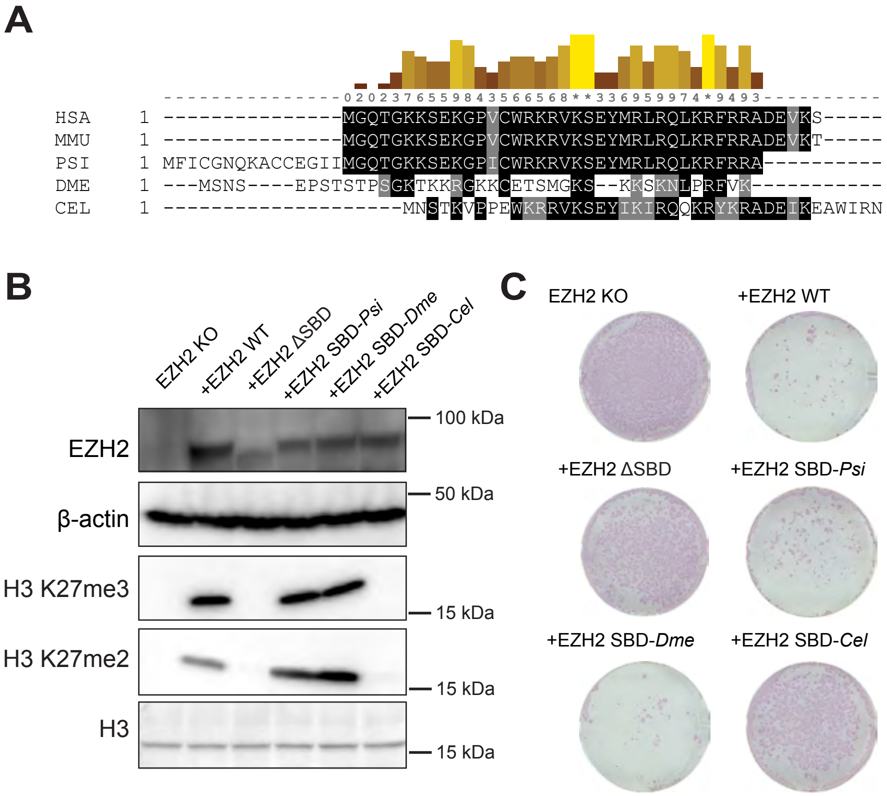


### SUPPLEMENTARY FIGURE 1

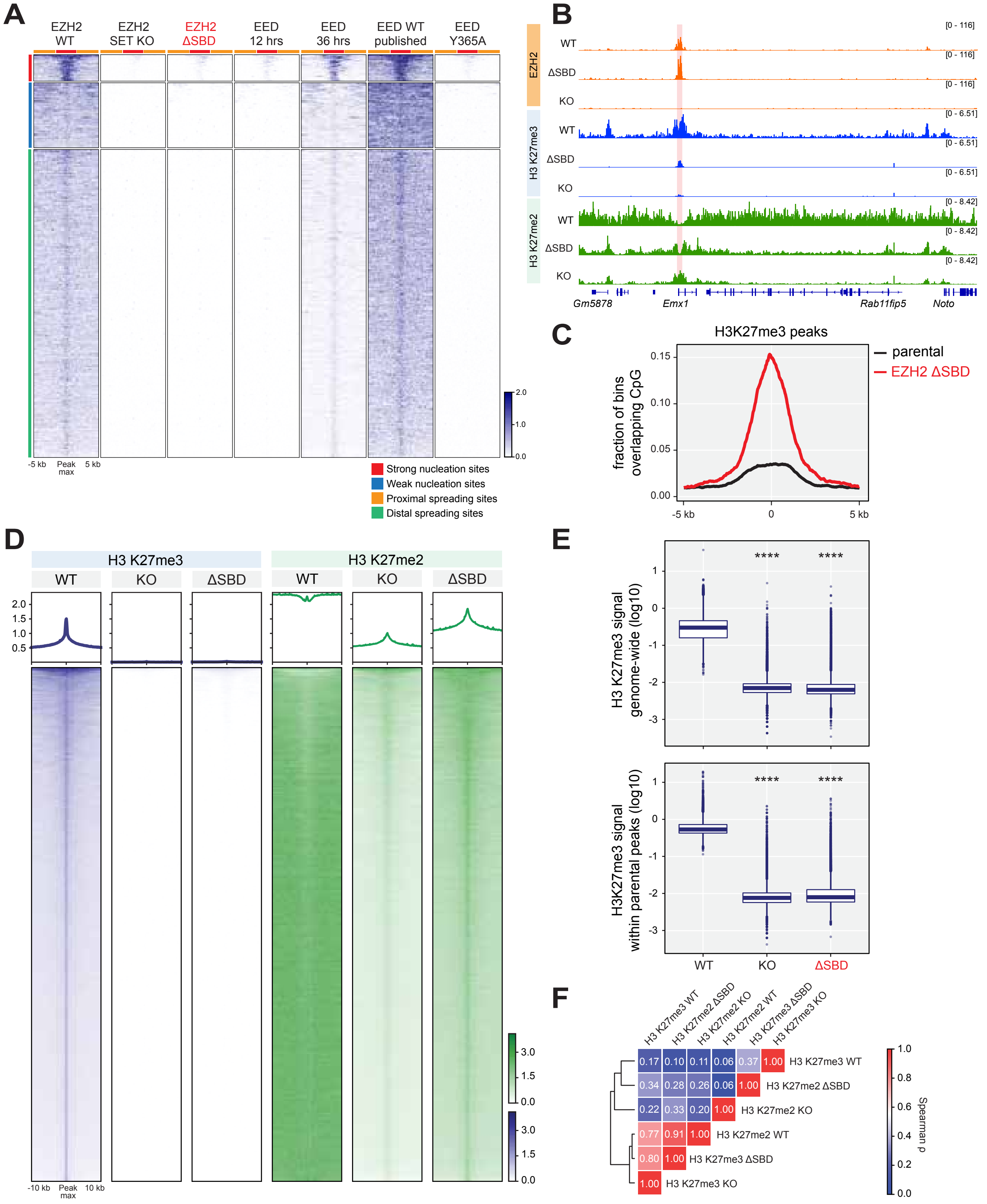


### SUPPLEMENTARY FIGURE 2

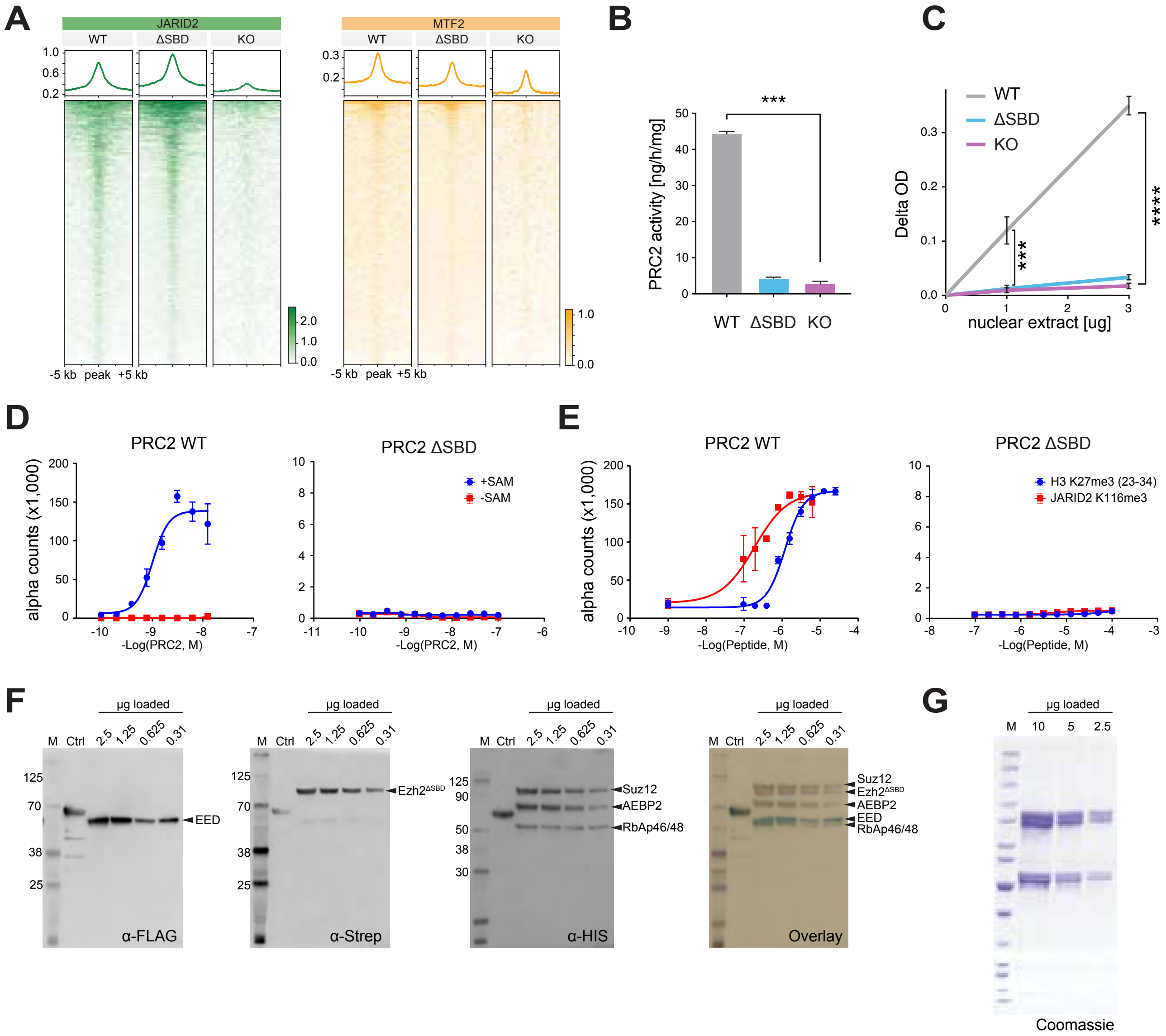


### SUPPLEMENTARY FIGURE 3

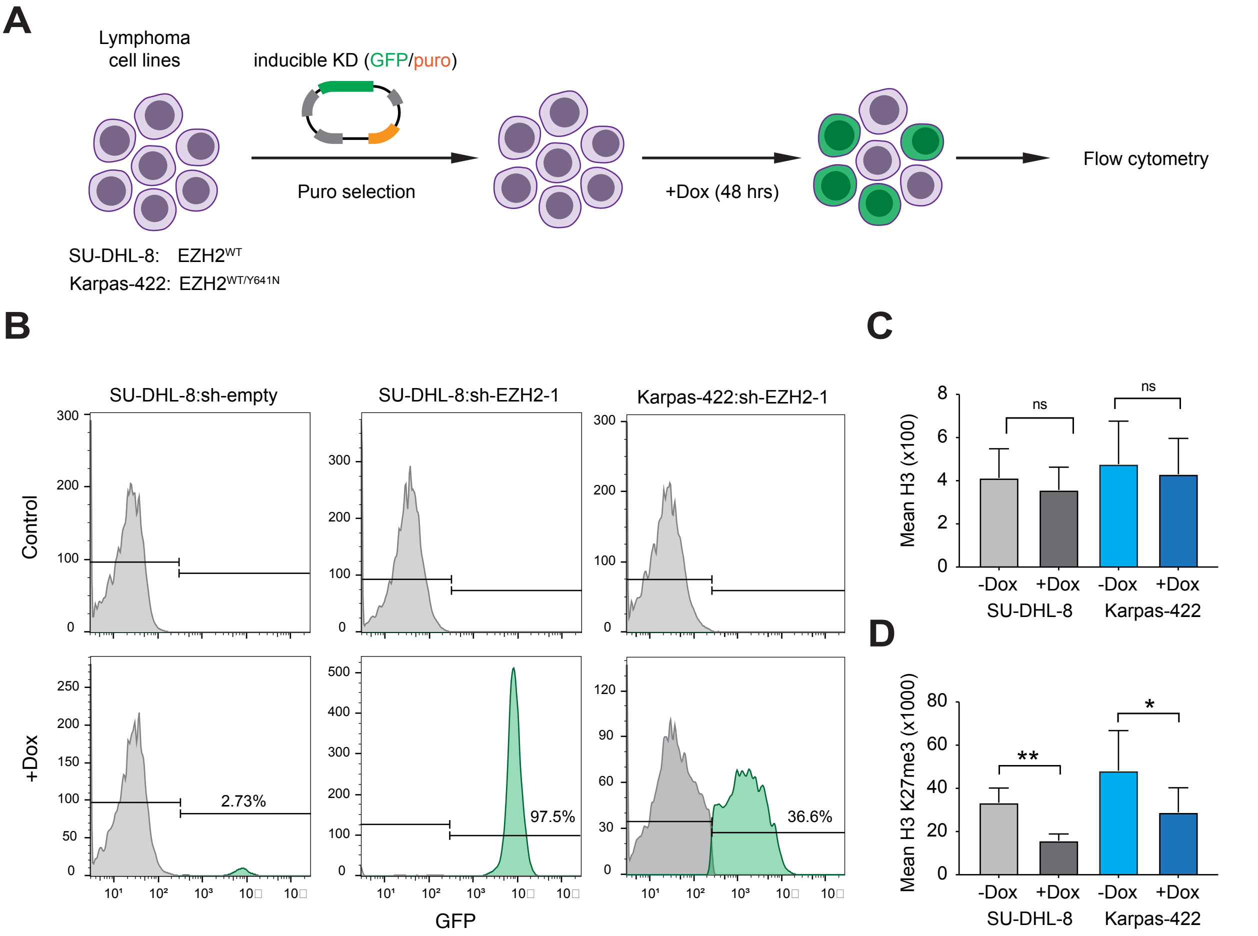


### SUPPLEMENTARY FIGURE 4

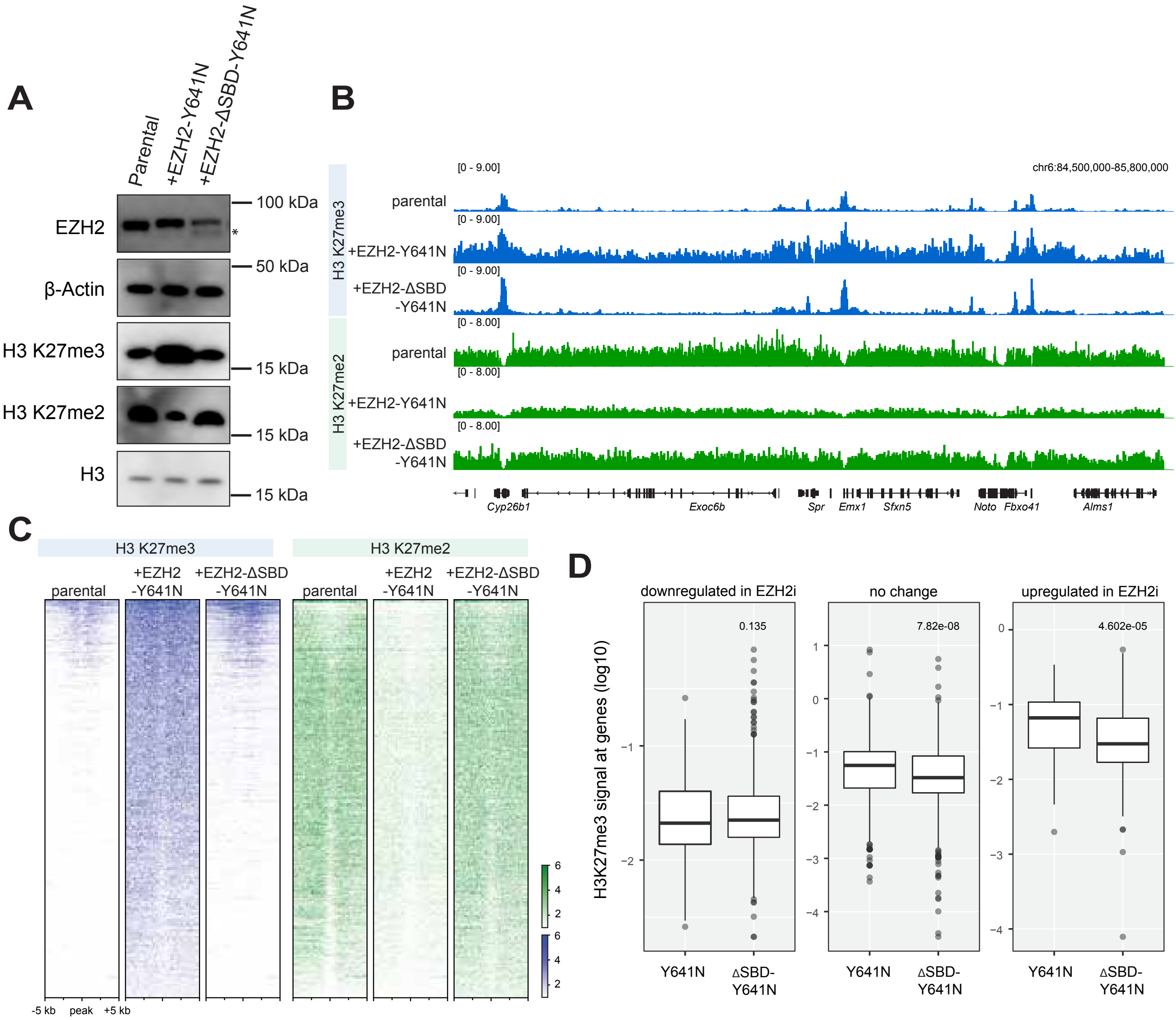


SUPPLEMENTARY FIGURE 5
